# Metabolic pairing of aerobic and anaerobic production in a one-pot batch cultivation

**DOI:** 10.1101/260216

**Authors:** Milla Salmela, Tapio Lehtinen, Elena Efimova, Suvi Santala, Rahul Mangayil

## Abstract

**Background:** The versatility of microbial metabolic pathways enables their utilization in vast number of applications. However, the electron and carbon recovery rates, essentially constrained by limitations of cell energetics, are often too low in terms of process feasibility. Cocultivation of divergent microbial species in a single process broadens the metabolic landscape and thus, the possibilities for more complete carbon and energy utilization.

**Results:** In this study, we integrated the metabolisms of two bacteria, an obligate anaerobe *Clostridium butyricum* and an obligate aerobe *Acinetobacter baylyi* ADP1. In the process, a glucose-negative mutant of *A. baylyi* ADP1 first deoxidized the culture allowing *C. butyricum* to grow and produce hydrogen from glucose. In the next phase, ADP1 produced long chain alkyl esters utilizing the by-products of *C. butyricum*, namely acetate and butyrate.

**Conclusions:** The cocultivation of strictly anaerobic and aerobic bacteria allowed the production of both hydrogen gas and long-chain alkyl esters in a simple one-pot batch process. The study demonstrates the potential of ‘metabolic pairing’ using designed microbial consortia for optimal electron and carbon recovery.

## Introduction

Almost all known biochemical reactions can be found in microbes, making them the most metabolically diverse class of organisms in the world. The natural versatility of microbes, and on the other hand, the development of cell engineering technologies have enabled their utilization in a great number of applications ranging from bioremediation of xenobiotics to the production of complex drug precursors [1-3]. Despite the advancements in cell-based processes enabled by metabolic engineering and synthetic biology, the laws of thermodynamics limit the number and efficiency of different metabolic pathways in a single organism. For example, the anaerobic fermentation of sugars to short-chain alcohols or organic acids is energetically favourable and redox balanced for cells, which makes the process scale-up rather straight forward and economically feasible. As a trade-off, however, in in these catabolic “downhill” pathways the substrate is converted to a product with lower energy and carbon content, causing substantial but inevitable loss of carbon. In contrast, the synthesis of many industrially relevant products with long carbon chain, such as alkanes, fatty acids, and alkyl esters, is energetically expensive for cells, and thus harnessing such thermodynamically “uphill” synthesis pathways for a profitable process is challenging. Recent advances in exploiting microbial electrosynthesis (MES) hold great potential for the energy recovery by external electron transfer, but further efforts are required to overcome the challenges related to the limitations of metabolic capabilities and systematic strain engineering for greater efficiency [4-5].

A potential strategy for combining a broader range of metabolic attributes is to utilize two or more different species or strains in a single process. In recent years, growing attention has been given to rationally designed microbial consortia addressing the issues related to the constraints of a single cell, such as limited substrate range or biosynthetic efficiency. For example, Hanly and Henson (2013) modelled substrate conversion by simultaneous utilization of hexoses and pentoses in a cocultivation by a respiratory deficient *Saccharomyces cerevisiae* and *Scheffersomyces stipitis* [6]. In another example, Minty et al. (2013) demonstrated the production of isobutanol directly from lignocellulose hydrolysate by a coculture of a fungi *Trichoderma reesei* and a bacterium *Escherichia coli.* In the culture, *T. reesei* secreted cellulases to create solubilized sugars, which were utilized by *E. coli* and further converted to isobutanol [7]. Park et al. (2012) also used cellulase-excreting fungi in a coculture. In their study, *Acremonium cellulolyticus* was used in a one-pot approach together with *Saccharomyces cerevisiae* to produce ethanol [8]. Similarly, Wang et al. (2015) used a bacterium *Clostridium celevecrescens* together with *Clostridium acetobutylicum* ATCC824 to produce butanol [9].Zhang et al. (2015), on the other hand, reported a coculture system, which involved a partial distribution of the metabolic pathway for cis-cis-muconic acid production in two engineered *E. coli* strains [10]. As a result of efficient intermediate redirection and alleviated burden to cells, the production of cis-cis-muconic acid was significantly improved.

Other typical issues related to single-cell cultures include excessive byproduct formation and poor tolerance against certain feedstock components and/or the (by)product itself. Organic acids, such as acetate, impose a great challenge in bioprocesses both in terms of toxicity to cells and as a drain for carbon and energy [8]. We have previously demonstrated the benefits of acetate redirection to biomass and product in a consortium of *E. coli* and *Acinetobacter baylyi* ADP1 [12]. In the culture, engineered *A. baylyi* ADP1 consumed the acetate produced by *E. coli*, which improved the biomass production and recombinant protein expression compared to single-cell cultures. More recently, we have successfully combined MES with long-chain ester synthesis by sequentially culturing an acetogenic strain *Sporumusa ovata* and *A. baylyi* ADP1 [13]. In the separated two-stage process, the acetogenic strain utilized carbon dioxide and electricity to produce acetate, which was subsequently fed to ADP1 to produce long-chain alkyl esters. In a somewhat reverse culture setup, we established a coculture of *A. baylyi* and *Clostridium butyricum* for the production of hydrogen (H2) gas from simulated and rice straw hydrolysates [14].

The glucose-negative *A. baylyi* selectively removed the *C. butyricum* growth inhibitors, namely acetate, formate, 4-hydroxybenzoate, and oxygen from culture, allowing the subsequent growth and H2 production by *C. butyricum* from hydrolysate sugars.

In the present study, we demonstrate the successful integration of anaerobic fermentation (representing the ‘downhill’ pathway) to aerobic synthesis (the ‘uphill’ pathway). In the process of two metabolically diverse bacterial species, aerobic and anaerobic phases alternate allowing a complete conversion of carbon and electrons to biomass and products in a non-optimized simple batch culture. As an ultimate example of a ‘carbon wasting’ downhill process, glucose is fermented to produce H2 by the obligate anaerobe *C. butyricum.* The byproducts, namely acetate and butyrate, together with naturally occurring oxygen, are converted to highly energy and carbon rich alkyl esters with an average of 34-carbon chains by the strict aerobe *A. baylyi* ADP1. This study demonstrates the potential of combining distinctive bacterial metabolisms for optimal electron and carbon recovery.

## Results

In order to allow efficient distribution and comprehensive consumption of carbon in a cocultivation of *C. butyricum* and *A. baylyi*, a previously engineered knock-out strain of *A. baylyi* deficient in glucose consumption was deployed [14]. The strain *A. baylyi* ADP1 Δgcd, designated here as ADP1-g, lacks the gene for glucose dehydrogenase, which is responsible for the first reaction in the modified Entner–Doudoroff pathway of *A. baylyi* sugar catabolism [15]. We hypothesized that in the cocultivation *C. butyricum* consumes the glucose and produces H2, followed by the production of long chain alkyl esters by ADP1-g from the liquid end-metabolites (acetate and butyrate) of *C. butyricum* (Fig. 1). In the first stage of the process, the growth medium is deoxidized by the metabolic activity of ADP1-g strain, allowing the growth and H2 production of the obligate aerobe *C. butyricum.* After C. butyricum growth, oxygen is allowed to enter the system, enabling ADP1-g growth and wax ester production from the residual carbon, namely acetate and butyrate.

**Figure 1.**
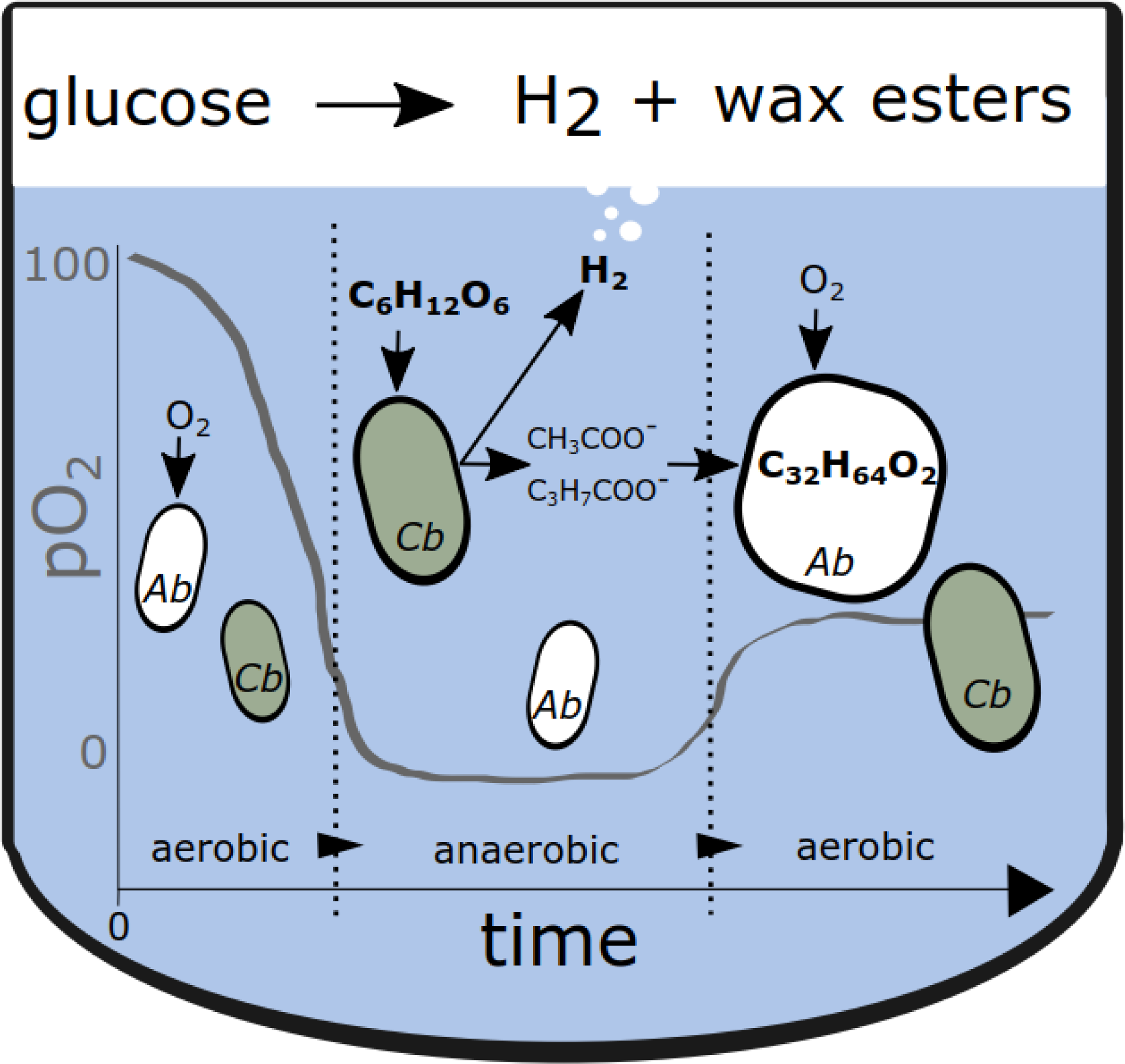
Schematic representation of the one-pot batch cultivation of *C. butyricum* and ADP1-g strain. In the first stage, the metabolic activity of ADP1-g deoxidizes the culture, allowing H2 production from glucose by *C. butyricum*. Thereafter, oxygen is released to the culture, enabling wax ester production from the residual carbon, namely acetate and butyrate, by ADP1-g.

### Suitability of ADP1-g for the alternating bioprocess conditions

We investigated the capability of the obligate aerobe ADP1-g to survive and subsequently grow in a cultivation with varying oxygen availability. In this experiment, ADP1-g was cultivated with limited oxygen supply in sealed vessels. The cells grew for 30 hours, after which the growth ceased due to oxygen depletion (Fig. 2). These anoxic growth conditions were maintained for another 24 hours and then oxygen was released to the vessels. Shortly after, an exponential increase in cell density was observed demonstrating that ADP1-g maintains cell viability after exposure to oxygen-deprived conditions for 24 hours.

**Figure 2.**
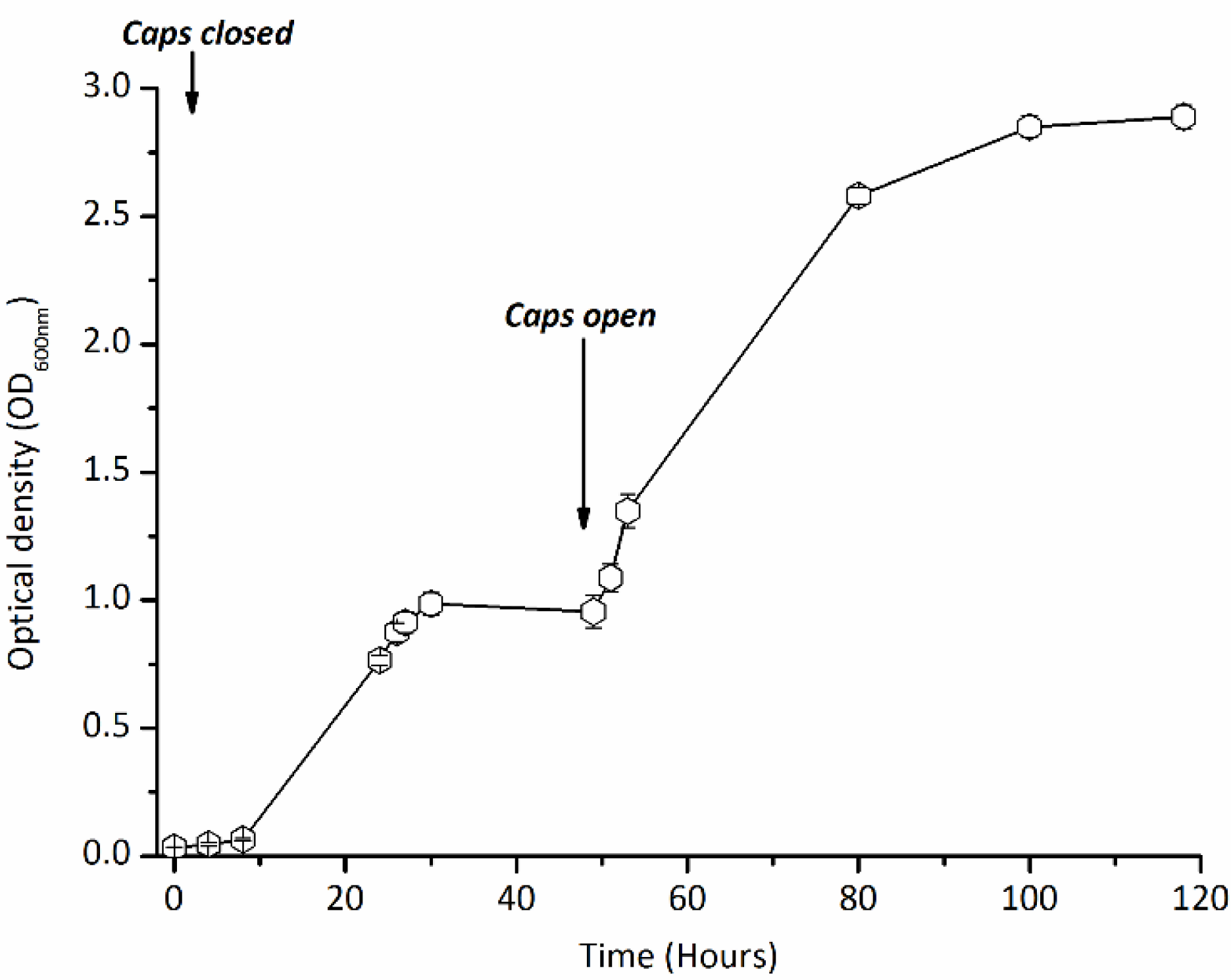
ADP1-g growth during alternating oxygen availability. The strain ADP1-g ability to withstand anoxic environment was conducted in 10 ml sterile aerobic MA/9 medium amended with 0.2% casein amino acids. Upon inoculation, the cultivation tubes were closed until growth cessation (30 hours) and opened at 50 hour time-point to resume the growth. The data points and error bars represent mean value and standard deviation from triplicate experimental repeats, respectively. In some cases, the error bars are smaller than the symbol.

Another growth test was conducted with ADP1-g to investigate the utilization of acetate and butyrate, which are potential end-products of *C. butyricum* in the cocultivation system. The strain consumed both carbon sources (initial 15 mM butyrate and 25 mM acetate) within 21 hours (Fig 3a.). In addition, wax esters were produced and accumulated up to 18 hours of cultivation (Fig 3b). Thereafter, the amount of WEs decreased due to the carbon depletion, which is a typical phenomenon in carbon limiting conditions [16].

**Figure 3.**
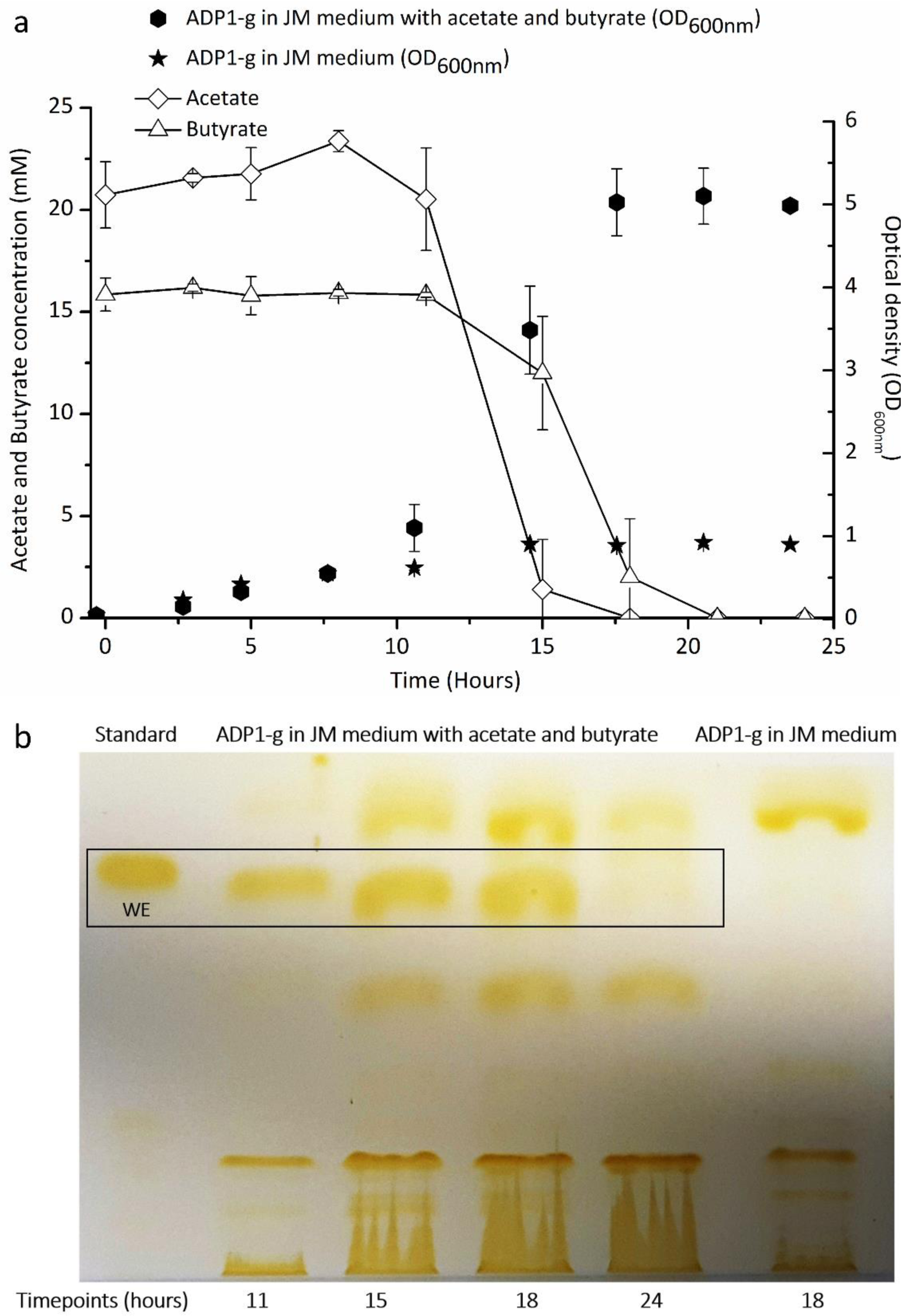
Growth, substrate utilization and wax ester production of ADP1-g cultivated in JM medium with 20mM acetate and 15mM butyrate. (a) The growth (closed symbols), determined as optical density at 600 nm wavelength, and organic acid utilization (open symbols). The presented data points are averaged values and error bars indicate standard deviation from triplicate experimental repeats. In some cases, the error bars are smaller than the symbol. (b) Thin layer chromatography analysis of wax ester synthesized by ADP1-g from acetate and butyrate. The time point samples analyzed with TLC correspond to the same presented in (a). In above figures a and b, ADP1-g cells in JM medium represents the growth and wax ester profiles, respectively, of ADP1-g cells when cultivated without acetate and butyrate supplementation (substrate blank).

### Cocultivation of ADP1-g and *C. butyricum* in batch

In the next stage, ADP1-g and *C. butyricum* were cocultivated in a 300 ml batch culture. In this one-pot approach, aerobic and anaerobic phases alternate allowing both bacteria to grow. Anoxic conditions were produced and maintained by the metabolic activity of the aerobic ADP1-g. The metabolite formation by *C. butyricum*, as well as the growth and end-products of both *C. butyricum* and ADP1-g, were determined. Twenty millimolar of glucose was used as carbon source for *C. Butyricum* growth based on preliminary tests conducted with different substrate concentrations (Additional file 1: Table S1). Ten millimolar acetate supplementation was used for initial growth and deoxygenation by ADP1-g according to ADP1-g acetate utilization trend (Table 1).

**Table 1.** Growth, pH and carbon utilization trend of ADP1-g cultivated in JM medium supplemented with glucose and acetate.

| Cultivation time (hours) <sup>a</sup> | Growth (OD<sub>600nm</sub>) <sup>b</sup> | Medium pH <sup>b</sup> | Glucose concentration (mM) <sup>b</sup> | Acetate concentration (mM) <sup>b</sup> |
| --- | --- | --- | --- | --- |
| 0 | 0.1±0.0 | 7.8±0.0 | 17.2±0.2 | 8.5±0.1 |
| 24 | 0.7±0.0 | 7.3±0.0 | 17.1±0.4 | BD |
| 27 | 1.1±0.1 | 7.9±0.1 | 17.6±1.4 | BD |
| 44 | 1.0±0.1 | 8.5±0.1 | 17.4±0.3 | BD |

<sup>a</sup> Sampling time similar as indicated in Figure 4
<sup>b</sup> Averaged data from triplicate experimental repeats ± standard deviation
BD, below detection limit

During the anaerobic phase, *C. butyricum* produced 1.6±0.1 mol H2/mol glucose with a cumulative H2 production of 131.5±5.2 ml. A drop in pH associated with *C. butyricum* growth occurred (pH 5) and most of the supplemented glucose was consumed (Fig. 4 a-b). Additionally, 11.4±0.3 mM of butyrate and 7.7±0.1 mM of acetate were produced from the *C. butyricum* fermentation (Fig. 4b).

**Figure 4.**
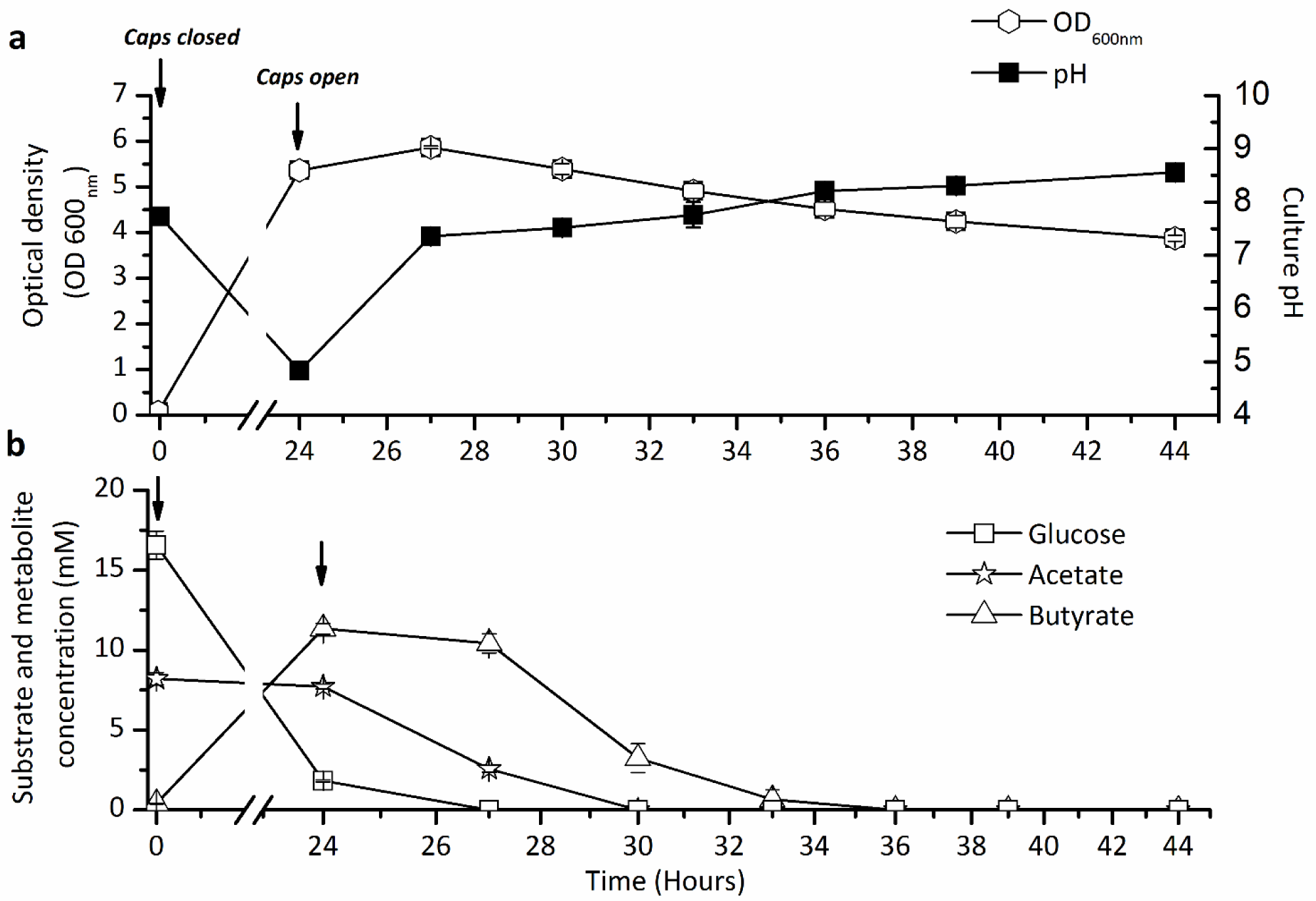
The growth, pH, and substrate concentrations of ADP1-g – *C. butyricum* cocultivations. Arrows indicate the time when the growth vessel caps were closed and opened during the experiment. (a) OD_600nm_ and pH trends of one-pot batch ADP1-g – *C. butyricum* cocultivations. (a) Glucose utilization (*C. butyricum*) and acetate-butyrate metabolism (ADP1-g) data from cocultivation experiment. The data points are averaged from triplicate experimental repeats. In some cases, the symbols overlap and the error bars (standard deviation) are smaller than the symbol.

After the *C. butyricum* fermentation, oxygen was released to the system enabling ADP1-g growth. Wax esters, the end products of ADP1, accumulated during 24-30 h of cultivations (Fig. 5). During this time, acetate was completely consumed along with most of the butyrate (Fig. 4b). The WEs were solely produced by ADP1-g, as *C. butyricum* does not produce WEs (Additional file 2: Fig. S2).

**Figure 5.**
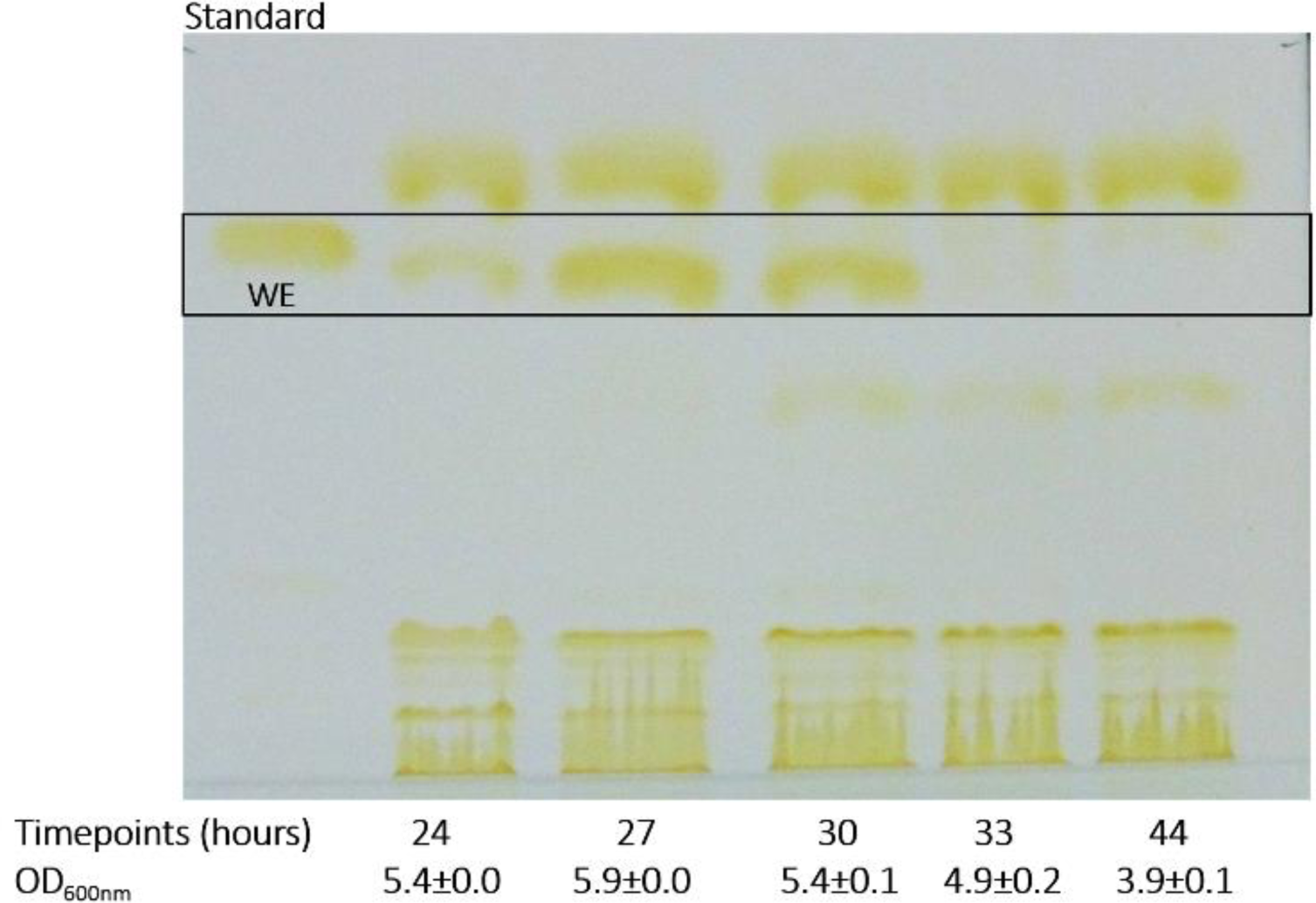
TLC analyses demonstrating the wax ester production from ADP1-g – *C. butyricum* coculture. The wax esters were synthesized by ADP1-g strain utilizing the acetate and butyrate generated during *C. butyricum* fermentation. The samples are from the same timepoints shown in Figure 4.

### Cocultivation of ADP1-g and *C. butyricum* in a 1-liter bioreactor

For a more detailed analysis of the coculture, a cultivation experiment was conducted in a 1L bioreactor. Similarly to the previous batch cultivation, ADP1-g and *C. butyricum* were simultaneously inoculated in the culture and the cultivation vessel was sealed. Oxygen partial pressure, pH, substrate/metabolite concentrations, and cell growth were monitored during the cultivation. The culture was deoxygenated by ADP1-g within 5 minutes after the inoculation (Fig. 6a) and anoxic conditions were maintained for the next 24 hours. During this anaerobic phase, *C. butyricum* consumed most of the supplemented glucose (83%) producing 4.8 mM of acetate, 8.3 mM of butyrate and 461 ml of H2 (cumulative). The carbon and electrons from the supplemented glucose were completely recovered in the metabolites and end products of *C. butyricum* (Additional file 3: Table S2). Within 3 hours of the latter aerobic phase, an increase in cell dry weight was observed as ADP1-g strain started to grow. During this aerobic stage, the strain recovered the carbon from *C. butyricum* metabolites for cell growth and storage compounds, namely wax esters (Figure. 6a and 6b, Table 2). Acetate was consumed rapidly in 4.5 hours whereas butyrate was consumed more gradually (Fig. 6b). The highest wax ester yield (30 mg/g biomass) was achieved after 8 hours of aerobic cultivation (32 hours of overall cultivation) with 100% of acetate and 61 % of butyrate consumed. The total yields of products recovered from the batch cocultivation were 1.7 mol H2 /mol glucoseconsumed and 10.8 mg WE/g glucoseconsumed, respectively.

**Table 2.**
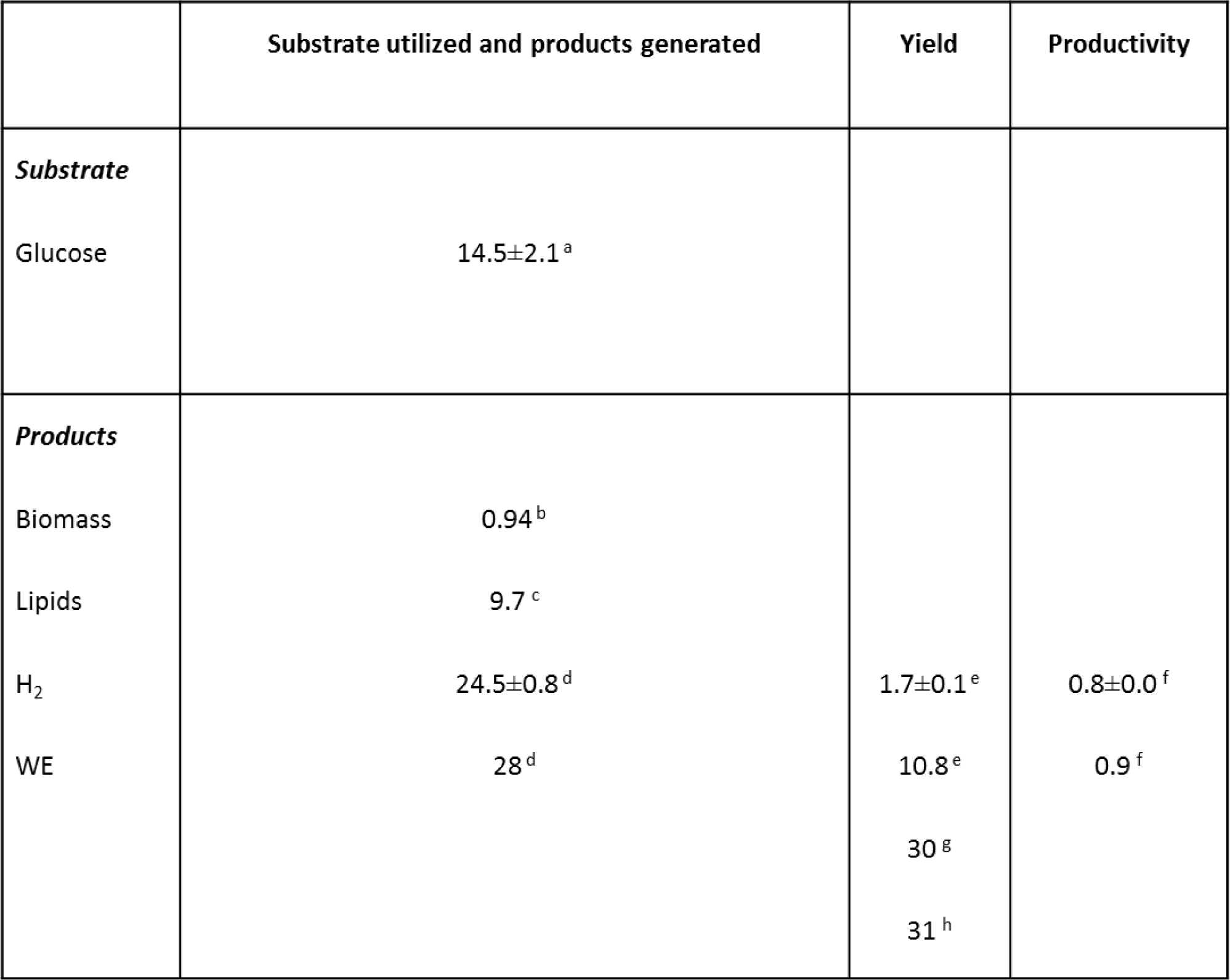
Products generated from glucose in one-pot batch cultivation by *C. butyricum* and ADP1-g.

**Figure 6.**
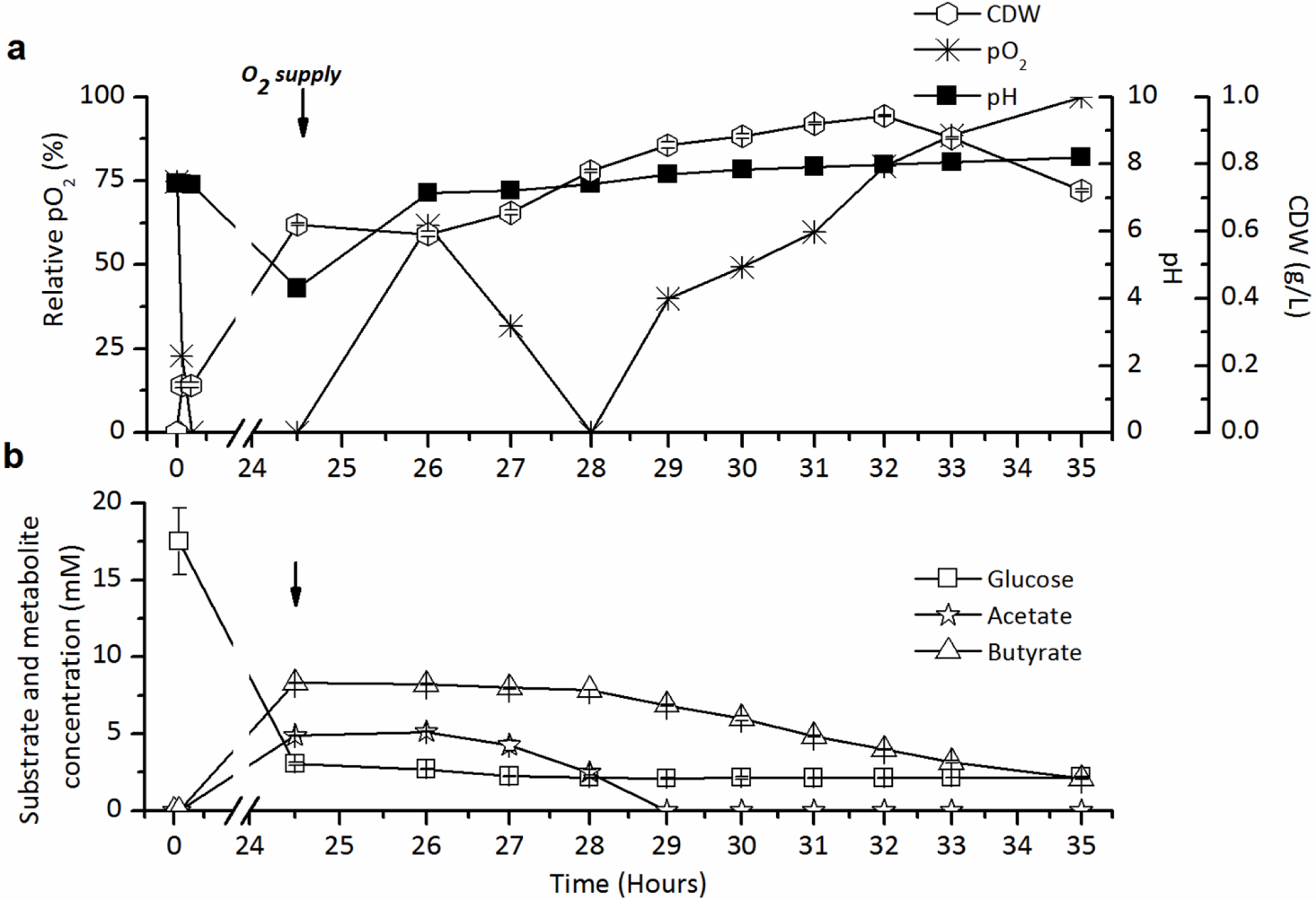
One-pot batch cultivation of ADP1-g and *C. butyricum* in 1-litre bioreactor. The cocultivation was carried out in 800 ml JM medium supplemented with 20 mM glucose. The changes in (a) pH, pO2 and biomass formation (in cell dry weight, CDW), and (b) glucose, acetate and butyrate concentrations are presented. *C. butyricum* consumed glucose producing acetate and butyrate during the first 24 hours, whereas ADP1-g utilized the fermentation end-products during the following 10 hours. Arrows indicate the timepoint at which the oxygen supply was initiated and the culture pH adjusted to 7.3. The data for CDW, glucose, acetate and butyrate are mean values from triplicate technical repeats. The standard deviations (error bars) of each data are plotted and in some cases the symbols overlap the error bars.

## Discussion

Efficient recovery of carbon and electrons in the final product is crucial in biological production processes. For cells, however, the optimal balance between carbon utilization and redox state often requires the by-production of unwanted metabolites. Moreover, the product range may be constrained depending on the metabolic capabilities of the cell. To address these issues, we designed and constructed a ‘metabolic pair’ of anaerobic and aerobic metabolisms enabling both fermentative hydrogen production and aerobic lipid synthesis with complete carbon and energy recovery in a simple one-pot batch cultivation. In the coculture set-up, a strict anaerobe *C. butyricum* was deployed for the production of H2 gas using glucose as the source of carbon and energy, whereas the strictly aerobic ADP1-g was utilized for maintaining oxygen-free conditions during the anaerobic fermentation, followed by the recovery of any residual carbon in the culture for the aerobic production of wax esters. In the batch process carried out in “one-pot”, aerobic and anaerobic phases alternate allowing toggle-like growth of the bacteria.

For the coculture system to be functional, the aerobic strain is required to both survive and sustain anaerobic conditions during the H2 production phase. Previously, ADP1-g has been shown in a proof of concept experiment to be able to deoxygenize growth media for *C. butyricum* [14], albeit the viability of the strain to survive anoxic conditions was not investigated. Thus, in our first experiment, the ability of ADP1-g to restore growth after anoxic phase was studied. According to the growth data, we found out that the strain was able to continue growth after being incubated in oxygen-limited conditions for 24 hours. In the subsequent experiment, we also confirmed that the strain is able to uphold sufficient anaerobic conditions during the oxygen-sensitive *C. butyricum* growth and H2 production phase. Examples of other approaches for employing oxygen-consuming bacteria as deoxygenizing agents have been previously reported. For example, cocultures for maintaining anoxic conditions (in a culture initially made anaerobic) have been made with *Enterococcus* [17]. Other cocultivation approaches have included systems to remove oxygen for *C. butylicum* growth with the help of *Bacillus subtilis* in order to increase butanol yields [18] [19]. Also, examples of deoxygenation of hydrogen producing cultures involving *C. butyricum* have been previously reported [17]. However, as the aerobic species do not contribute to product formation in these processes, the substrate utilized for their metabolism is “wasted” and lowers the yield of the overall process. In contrast, in our system, ADP1-g not only consumes the oxygen, but also scavenges and upgrades the otherwise wasted organic acids produced by *C. butyricum.* Notably, ADP1-g does not consume glucose, and thereby does not impair the efficiency of H2 production from glucose.

The incapability of ADP1-g to utilize glucose has been previously demonstrated [14]. Wax ester production by *A. baylyi* ADP1 using acetate as a carbon source has been also previously reported [20] whereas WE production from for butyrate has not been recorded to our knowledge. In our study, ADP1-g was shown to grow and produce alkyl esters (WEs) from both acetate and butyrate, which makes it ideal for upgrading the metabolites of *C. butyricum.* It is also notable that the strain was able to continue growth and establish wax ester production directly in the medium ‘used’ and occupied by *C. butyricum. C. butyricum* ferments glucose and other carbohydrates readily to H2 and volatile fatty acids (VFA), such as acetate and butyrate [21] [22]. In theory, the complete oxidation of glucose yields 12 mol of H2, though in reality, thermodynamic and metabolic restrictions in biological systems divert the substrate also towards biomass and other metabolites [23]. Thus, the theoretical yields of the NADH dependent H2 production by *Clostridium* vary between 2 – 4 mol-H2/mol-glucose depending on the ratio of produced metabolites butyrate and acetate. As the latter is associated with higher H2 yields - and *butyricum* typically favors butyrate production -the actual yields obtained are closer to the lower range of the theoretical values. In our experiments, the cocultivation yielded 1.7±0.1 mol H2/mol glucoseconsumed with an acetate to butyrate ratio of 0.6:1. As comparison, experiments with pure cultures of *C. butyricum* in optimized production conditions have yielded results such as 2.29 mol H2/mol glucose [22] and 2.21 mol H2/mol glucose [21] whereas coculture designs have yielded values such as 1.65 mol H2/mol glucose [24]. The metabolites (acetate and butyrate) produced in our experiments account up to 49,5% of the carbon and 59.1% of the electrons from the initial glucose supplementation, which are an untapped resource when using pure cultures. In our study, these compounds were scavenged in the second aerobic stage of the one-pot system providing additional value for the process in the form of 10.8 mg WE / g glucose_consumed_.

Co-culture systems allow the design of production systems with metabolic capabilities exceeding those of any individual organism. Zhou et al [25] produced oxygenated taxanes by dividing the complex pathway between *E. coli* and *S. cerevisiae. E. coli* produced taxadiene, which was oxygenated by *S. cerevisiae*, resulting in the production of complex oxygenated taxane molecules that have not been produced by any single microbe. In another example, Liu et al [26] used a consortium consisting of three different microbes for the division of labor in order to enhance the electricity production from glucose. In the consortium, *E. coli* first converted glucose to lactate, which was oxidized by *Shewanella oneidensis* to acetate. The acetate was further utilized by *E. coli. B. subtilis*, in turn, produced riboflavin, which is required as an electron shuttle. These studies exemplify how a single biosynthetic pathway can be divided between different organisms in order to improve the production. In the present study, we combine two very different metabolisms (aerobic and anaerobic, catabolic and anabolic) that complement each other for the efficient recovery of electrons (H2 and WE) and carbon (WE).

The relevance of the current platform could be further improved by engineering the production efficiency and/or product range. *A. baylyi* ADP1 can be readily engineered for improved production of native or non-native products [20, 27-29], whereas limited number of studies describing successful engineering of *C. butyricum* has been reported. Thus, one of the advantages of metabolically diverse cocultures is the modularity and flexibility of the system; substrate and product range can be individually engineered and tuned for applicable strain(s). Such division of engineering load lowers the metabolic burden in one strain, and on the other hand, reduces the challenges faced with strains for which engineering tools and methods are not readily available.

## Conclusions

Cocultivation of organisms with divergent metabolic characteristics can greatly broaden the substrate and product range and facilitates more comprehensive utilization of carbon and energy. We successfully integrated bacterial anaerobic and aerobic metabolisms in a synthetic consortium, and by exploiting the advantageous properties of both metabolisms, demonstrated the production of hydrogen gas and long-chain alkyl esters in a single process. The study demonstrates the power of metabolic pairing in resolving metabolic and thermodynamic limitations of single organisms as well as in developing novel metabolic combinations for more efficient carbon and energy recovery.

## Methods

### Strains and cultivation medium

*Acinetobacter baylyi* ADP1 (DSM 24193) devoid of glucose dehydrogenase gene ACIAD2983, (referred here as ADP1-g) and *Clostridium butyricum*, isolated from a hydrogen-producing bioreactor, were used in the study [30]. Unless otherwise indicated, ADP1-g cells cultivated in low-salt Lysogeny broth (LB) (tryptone, 10 g/liter; yeast extract, 5 g/liter; sodium chloride, 1 g/liter) at 30°C and 300rpm, were used as pre-inoculums. The growth kinetics of ADP1-g in a three stage bioprocess was studied in in MA/9 medium (g/L; 5.518 Na2HPO_4_·2H_2_O, 3.402 KH_2_PO_4_, 1 NH_4_Cl, 0.008 nitrilotriacetic acid, 0.487 FeCl_3_, 0.25 MgSO_4_·7H_2_O, 0.02 CaCl_2_·2H_2_O, 0.2% casein amino acids and 2ml/L of SL7 trace element solution). ADP1-g acetate and butyrate utilization experiments were conducted in a minimal medium [24] (referred as JM medium) with modifications as described in [14]. *C. butyricum* precultivations were conducted in anaerobic Reinforced Clostridial Medium (RCM, Sigma Aldrich, US) at 37°C and 150rpm. The co-cultivation experiments, using ADP1-g and C. *butyricum*, were conducted in JM medium and the cultures were grown at 30°C and 300rpm.

### Experimental procedure

The growth test to investigate ADP1-g viability in the three stage aerobic-anaerobic-aerobic bioprocess was conducted in tubes containing 10 ml of sterile aerobic MA/9 medium supplemented with casein amino acids. Upon inoculating (initial optical density 0.02), the cultivation tubes were closed with sterile septum rubber stoppers, tightened with aluminum crimps and incubated at 30°C and 300rpm. A blank cultivation (devoid of inoculant) was included as control to determine ‘false positive’ results from contamination and as analytical control. The experiment was conducted in triplicates and the cell growth data measured at specific intervals using a spectrophotometer (Ultrospec 500 pro, Amersham Biosciences, UK) for 120 hours, were averaged.

Batch experiments to investigate ADP1-g capacity to utilize organic acids from *C. butyricum* fermentation was performed in 120 ml serum bottles with a working volume of 50 ml sterile JM medium. Acetate and butyrate were supplemented at concentrations 20 mM and 15 mM, respectively. The inoculated cells (initial cell density 0.02) were cultured aerobically at 30°C and 300rpm. The cells were also inoculated to a substrate blank, i.e. cultivation medium devoid of acetate and butyrate. Samples to analyze cell growth, organic acid utilization, lipid analysis and medium pH were collected in 3-hour interval for 24 hours.

The co-cultivations of ADP1-g and *C. butyricum* were conducted in triplicate 500ml batch bottles containing 300ml of sterile aerobic JM medium. To initiate ADP1-g and *C. butyricum* growth, the medium was supplemented with 10 mM acetate and 20 mM glucose, respectively. Two milliliters of precultivated ADP1-g with an optical density at wavelength 600nm (OD_600nm_) of 3.2 was inoculated alone and coinoculated with 2 ml of *C. butyricum* precultivation (OD_600nm_ 3.1). ADP1-g cultivated in similar medium and a blank medium were used as the controls in this experiment. Following inoculation, the batch bottles were capped (rubber stoppers), tightened (aluminum crimps). Initial samples to determine cell growth, substrate utilization, liquid metabolites and lipids were collected and the cultures were incubated at 30°C and 200rpm. The gaseous end products were analyzed from the bottle headspace after a 24 hour cultivation period and the rubber stoppers were removed. Opening the bottle caps ensured the end of anaerobic phase and culture samples for analysis were collected. The medium pH, after the anaerobic phase, from the coculture, ADP1-g alone and blank cultivations were measured aseptically and adjusted to pH 7.4 (initial pH) with 5 M sterile sodium hydroxide solution. The cultures were divided to sterile 50 ml serum bottles and incubated under aerobic conditions at 30°C and 200rpm. The experiment was conducted in triplicates and samples were collected every 3 hours during 20-hour cultivation for analysis.

Bioreactor experiment was conducted in a sterile 1-liter vessel (Sartorius Biostat B plus Twin System, Germany) with an online pH monitoring system. To a medium volume of 795 ml, 50ml of ADP1-g precultivated in JM medium supplemented with 25 mM acetate (OD_600nm_ 3.9) and 5 ml of *C. butyricum* precultivated in RCM (OD_600nm_ 2.3) were inoculated, totaling the cultivation volume to 850 ml. To prevent gas leakage, the fermentor outlets were closed and a gas collection bag (Supelco, USA) was connected to the exhaust. After initial sample collection, the cultures were grown at 30°C and 350 rpm for 24 hours. At the end of anaerobic phase, the collection bag was removed, samples were collected for analysis and the medium pH was adjusted 7.3 with sterile 5 M NaOH solution. The aerobic phase was initiated with air supply. The medium pH and partial O2 (pO2) profiles were obtained from the bioreactor. From the bioreactor, 45 ml of culture was removed every hour from which duplicate technical repeats to monitor cell growth and organic acid utilization were collected and the remaining culture (~40ml) was used for quantitative lipid analysis.

### Analytical techniques

For qualitative lipid analysis, 4 ml of culture samples were analyzed using thin layer chromatography as described in [16]. For dry cell weight (DCW) calculations in bioreactor experiment, 40 ml of original cultures were pelleted in pre-weighed tubes at 30,000 g (4°C), freeze-dried and weighed. The tube weights were subtracted from the freeze-dried cell weights to determine the DCW (g/L). The lipids extracted from the freeze-dried cells were quantitatively analyzed using NMR as described in [28].

Glucose, acetate and butyrate concentrations were analyzed, in triplicates, using High-Performance Liquid Chromatography (HPLC) (LC-20AD, Shimadzu, Japan) equipped with equipped with Shodex SUGAR (SH1011) column (300 × 8 mm), refractive index detector (RID, RID-10A) and 0.01N H2SO4 as mobile phase. One milliliter of culture samples were centrifuged at 15,000 g for 7 minutes (4°C), filtered through 0μm polycarbonate filter (Chromafil^®^ PET-45/25, Macherey-Nagel, Germany) and diluted tenfold with ultrapure water in 1ml HPLC vials. The injection volume, column temperature and mobile phase flowrate was set to 100μ!, 40°C and 0. 6 ml/min, respectively. Identification and quantification of carbon substrate and liquid fermentation metabolites were based on chromatography using external standards.

A gas chromatograph (GC-2014, Shimadzu GC) fitted with a thermal conductivity detector, PORAPAK column (2 m × 2 mm) and N2 as carrier gas was used to analyze the gaseous end products as described in [31]. In brief, to the GC maintained with column, oven and detector temperatures of 80°C, 80°C and 110°C, respectively, 200μl of sample from the collection bag was injected to the sampling port. The overpressure, i.e. gas volume exceeding the headspace volume of the vessel, was analyzed using water displacement method. Hydrogen yields (mol H2 / mol glucoseconsumed) were calculated by converting the GC data (ml) and overpressure values (ml) to millimolar (mM) values using the ideal gas law equation (for room temperature) and dividing it with the product of cultivation volume (L) and utilized sugar (mM). For batch experiments, each experimental repeats were measured twice and in bioreactor experiment, triplicate technical repeats were included.

Carbon balances of *C. butyricum* gaseous and liquid metabolites and utilized substrate were calculated by multiplying the concentrations with the respective number of carbon in the molecular formula. The CO2 in the liquid phase was ignored during the carbon balance calculations. Electron mass balances were calculated by multiplying the carbon mass of utilized glucose and each metabolite with the corresponding degree of reduction (mol electrons per C-mol). For carbon and electron mass calculations, the chemical formula of *C. butyricum* biomass was assumed to be CH1.624O0.456N0.216P0.033S0.0047 [32]. The biomass concentrations (mM) after anaerobic cultivation was calculated by dividing the DCW with the total mass from the formula (g/mol). Both carbon and electron recoveries were calculated by determining the percentage of sum total of carbon and electron mass of liquid and gaseous metabolites divided with the respective masses calculated for the utilized sugar.

## Declarations

### Authors’ contributions

RM and SS designed the system and planned the experimental setup. MS, TL, and RM executed the experimental work. EE and TL carried out the lipid analytics. MS, TL, SS and RM interpreted the data and wrote the manuscript. RM supervised the work. All authors read and approved the final manuscript.

### Availability of data and materials

All the data analysed in this study are included in this manuscript and its additional files.

### Ethics approval and consent to participate

Not applicable

### Consent for publication

Not applicable

## Acknowledgements

Not applicable

## References

1. Dvořák P, Nikel PI, Damborský J, de Lorenzo V. Bioremediation 3.0: Engineering pollutant-removing bacteria in the times of systemic biology. Biotechnology Advances. 2017;35:845–66.

2. Paddon CJ, Westfall PJ, Pitera DJ, Benjamin K, Fisher K, McPhee D, et al. High-level semi-synthetic production of the potent antimalarial artemisinin. Nature. 2013;496:528–32.

3. Eudes A, Benites VT, Wang G, Baidoo EEK, Lee TS, Keasling JD, et al. Precursor-directed combinatorial biosynthesis of cinnamoyl, dihydrocinnamoyl, and benzoyl anthranilates in saccharomyces cerevisiae. PLoS One. 2015;10:e0138972.

4. Kracke F, Lai B, Yu S, Krömer JO. Balancing cellular redox metabolism in microbial electrosynthesis and electro fermentation – a chance for metabolic engineering. Metab Eng. 2018;45:109–20.

5. Teravest MA, Ajo-Franklin CM. Transforming exoelectrogens for biotechnology using synthetic biology. Biotechnol Bioeng. 2016;113:687–97.

6. Hanly TJ, Henson MA. Dynamic metabolic modeling of a microaerobic yeast coculture: predicting and optimizing ethanol production from glucose/xylose mixtures. Biotechnol Biofuels. 2013;6:44.

7. Minty JJ, Singer ME, Scholz SA, Bae C-H, Ahn J-H, Foster CE, et al. Design and characterization of synthetic fungal-bacterial consortia for direct production of isobutanol from cellulosic biomass. Proc Natl Acad Sci. 2013;110:14592–7.

8. Park EY, Naruse K, Kato T. One-pot bioethanol production from cellulose by coculture of *Acremonium cellulolyticus* and *Saccharomyces cerevisiae*. Biotechnol Biofuels. 2012;5:64.

9. Wang Z, Cao G, Zheng J, Fu D, Song J, Zhang J, et al. Developing a mesophilic coculture for direct conversion of cellulose to butanol in consolidated bioprocess. Biotechnol Biofuels. 2015;8:84.

10. Zhang H, Li Z, Pereira B, Stephanopoulos G. Engineering *E. coli-E. coli* cocultures for production of muconic acid from glycerol. Microb Cell Fact. 2015;14:134.

11. Luli GW, Strohl WR. Comparison of growth, acetate production, and acetate inhibition of *Escherichia coli* strains in batch and fed-batch fermentations. Appl Environ Microbiol. 1990;56:1004–11.

12. Santala S, Karp M, Santala V. Rationally engineered synthetic coculture for improved biomass and product formation. PLoS One. 2014;9:e113786.

13. Lehtinen T, Efimova E, Tremblay PL, Santala S, Zhang T, Santala V. Production of long chain alkyl esters from carbon dioxide and electricity by a two-stage bacterial process. Bioresour Technol. 2017;243:30–6̤

14. Kannisto MS, Mangayil RK, Shrivastava-Bhattacharya A, Pletschke BI, Karp MT, Santala VP. Metabolic engineering of *Acinetobacter baylyi* ADP1 for removal of *Clostridium butyricum* growth inhibitors produced from lignocellulosic hydrolysates. Biotechnol Biofuels. 2015;8:198.

15. Taylor WH, Juni E. Pathways for biosynthesis of a bacterial capsular polysaccharide. I. Carbohydrate metabolism and terminal oxidation mechanisms of a capsuleproducing coccus. J Bacteriol. 1961;81:694–703.

16. Santala S, Efimova E, Karp M, Santala V. Real-Time monitoring of intracellular wax ester metabolism. Microb Cell Fact. 2011;10:75.

17. Yokoi H, Tokushige T, Hirose J, Hayashi S, Takasaki Y. H2 production from starch by a mixed culture of *Clostridium butyricum* and *Enterobacter aerogenes*. Biotechnol Lett. 1998;20:143–7.

18. Tran HTM, Cheirsilp B, Hodgson B, Umsakul K. Potential use of *Bacillus subtilis* in a co-culture with *Clostridium butylicum* for acetone–butanol–ethanol production from cassava starch. Biochem Eng J. 2010;48:260–7.

19. Abd-Alla MH, Elsadek El-Enany A-W. Production of acetone-butanol-ethanol from spoilage date palm (Phoenix dactylifera L.) fruits by mixed culture *of Clostridium acetobutylicum* and *Bacillus subtilis*. Biomass and Bioenergy. 2012;42:172–8.

20. Kannisto M, Efimova E, Karp M, Santala V. Growth and wax ester production of an *Acinetobacter baylyi* ADP1 mutant deficient in exopolysaccharide capsule synthesis. J Ind Microbiol Biotechnol. 2017;44:99–105.

21. Chong M-L, Abdul Rahman N, Yee PL, Aziz SA, Rahim RA, Shirai Y, et al. Effects of pH, glucose and iron sulfate concentration on the yield of biohydrogen by *Clostridium butyricum* EB6. Int J Hydrogen Energy. 2009;34:8859–65.

22. Lin P-Y, Whang L-M, Wu Y-R, Ren W-J, Hsiao C-J, Li S-L, et al. Biological hydrogen production of the genus *Clostridium:* Metabolic study and mathematical model simulation. Int J Hydrogen Energy. 2007;32:1728–35.

23. Cai G, Jin B, Saint C, Monis P. Genetic manipulation of butyrate formation pathways in *Clostridium butyricum*. J Biotechnol. 2011;155:269–74.

24. Seppälä JJ, Puhakka JA, Yli-Harja O, Karp MT, Santala V. Fermentative hydrogen production by *Clostridium butyricum* and *Escherichia coli* in pure and cocultures. Int J Hydrogen Energy. 2011;36:10701–8.

25. Zhou K, Qiao K, Edgar S, Stephanopoulos G. Distributing a metabolic pathway among a microbial consortium enhances production of natural products. Nat Biotechnol. 2015;33:377–83.

26. Liu Y, Ding M, Ling W, Yang Y, Zhou X, Li BZ, et al. A three-species microbial consortium for power generation. 2017;10:1600–9.

27. Lehtinen T, Santala V, Santala S. Twin-layer biosensor for real-time monitoring of alkane metabolism. FEMS Microbiol Lett. 2017;364.

28. Santala S, Efimova E, Kivinen V, Larjo A, Aho T, Karp M, et al. Improved Triacylglycerol Production in *Acinetobacter baylyi* ADP1 by Metabolic Engineering. Microb Cell Fact. 2011;10:36.

29. Kannisto M, Aho T, Karp M, Santala V. Metabolic Engineering of *Acinetobacter baylyi* ADP1 for Improved Growth on Gluconate and Glucose. Appl Environ Microbiol. 2014;80:7021–7.

30. Koskinen PEP, Kaksonen AH, Puhakka JA. The relationship between instability of H2 production and compositions of bacterial communities within a dark fermentation fluidized-bed bioreactor. Biotechnol Bioeng. 2007;97:742–58.

31. Mangayil R, Santala V, Karp M. Fermentative hydrogen production from different sugars by *Citrobacter* sp. CMC-1 in batch culture. Int J Hydrogen Energy. 2011;36:15187–94.

32. Serrano-Bermúdez LM, González Barrios AF, Maranas CD, Montoya D. *Clostridium butyricum* maximizes growth while minimizing enzyme usage and ATP production: Metabolic flux distribution of a strain cultured in glycerol. BMC Syst Biol. 2017;11.

